# Signatures of selection at drug resistance loci in *Mycobacterium tuberculosis*

**DOI:** 10.1101/173229

**Authors:** Tatum D. Mortimer, Alexandra M. Weber, Caitlin S. Pepperell

## Abstract

Tuberculosis (TB) is the leading cause of death by an infectious disease, and global TB control efforts are increasingly threatened by drug resistance in *Mycobacterium tuberculosis (M. tb).* Unlike most bacteria, where lateral gene transfer is an important mechanism of resistance acquisition, resistant *M. tb* arises solely by *de novo* chromosomal mutation. Using whole genome sequencing data from two natural populations of *M. tb,* we characterized the population genetics of known drug resistance loci using measures of diversity, population differentiation, and convergent evolution.
We found resistant sub-populations to be less diverse than susceptible sub-populations, consistent with ongoing transmission of resistant *M. tb.* A subset of resistance genes (“sloppy targets”) were characterized by high diversity and multiple rare variants; we posit that a large genetic target for resistance and relaxation of purifying selection contribute to high diversity at these loci. For “tight targets” of selection, the path to resistance appeared narrower, evidenced by single favored mutations that arose numerous times on the phylogeny and segregated at markedly different frequencies in resistant and susceptible sub-populations. These results suggest that diverse genetic architectures underlie drug resistance in *M. tb,* and combined approaches are needed to identify causal mutations. Extrapolating from patterns observed in well-characterized genes, we identified novel candidate variants involved in resistance. The approach outlined here can be extended to identify resistance variants for new drugs, to investigate the genetic architecture of resistance, and, when phenotypic data are available, to find candidate genetic loci underlying other positively selected traits in clonal bacteria.

**Importance:** *Mycobacterium tuberculosis (M. tb),* the causative agent of tuberculosis (TB), is a significant burden on global health. Antibiotic treatment imposes strong selective pressure on *M. tb* populations. Identifying the mutations that cause drug resistance in *M. tb* is important for guiding TB treatment and halting the spread of drug resistance. Whole genome sequencing (WGS) of *M. tb* isolates can be used to identify novel mutations mediating drug resistance and to predict resistance patterns faster than traditional methods of drug susceptibility testing. We have used WGS from natural populations of drug resistant *M. tb* to characterize effects of selection for advantageous mutations on patterns of diversity at genes involved in drug resistance. The methods developed here can be used to identify novel advantageous mutations, including new resistance loci, in *M. tb* and other clonal pathogens.

## Introduction

*Mycobacterium tuberculosis (M. tb),* the causative agent of tuberculosis (TB), is estimated to have caused 1.4 million deaths in 2015, making it the leading cause of death due to an infectious disease. The proportion of TB due to MDR (multi-drug resistant) *M. tb* resistant to first line anti-tuberculosis drugs isoniazid (INH) and rifampin (RIF)) is increasing (1), which poses a major threat to global public health. Unlike most bacteria, *M. tb* evolves clonally, so resistance cannot be transferred among strains or acquired from other species of bacteria: drug resistance in *M. tb* results from *de novo* mutation within patients and transmission of drug resistant clones (2–4). The relative contributions of *de novo* emergence and transmitted drug resistance varies across sampling locations (5–9). Another potential variable in the emergence of drug resistant TB is *M.tb’s* lineage structure: seven distinct lineages have been identified among globally extant populations of *M. tb.* Among these, lineage 2 (L2) has been associated with relatively high rates of drug resistance, and it has been postulated that the acquisition of resistance is a result of higher rates of mutation in this lineage (10). Studies of *M.tb* evolution within hosts with TB have shown that emergence of drug resistance is associated with clonal replacements that lead to reductions in genetic diversity of the bacterial population (11, 12).

Many of the methods developed to identify advantageous mutations, such as those conferring antibiotic resistance, depend on recombination to differentiate target loci from neutral variants (13). However, in clonal organisms like *M. tb,* neutral and deleterious mutations that are linked to advantageous variants will evolve in tandem with them. This linkage among sites can also cause competition between genetic backgrounds with beneficial mutations, decreasing the rate of fixation for beneficial alleles, while deleterious alleles are purged less efficiently (14–16).

While the majority of the *M. tb* genome is subject to purifying selection (i.e. selection against deleterious mutations) (3), antibiotic pressure exerts strong selection for advantageous variants that confer resistance. *M. tb* drug resistance has been the focus of extensive investigation, and a variety of resistance mutations have been characterized for commonly used anti-tuberculosis drugs (17). Drug resistance mutations can be associated with fitness costs (18–20), and compensatory mutations that ameliorate these fitness costs have been identified in the context of rifampicin resistance (21, 22). Resistance mutations found to have lower fitness costs *in vitro* – as measured by competition assays – are found at higher frequencies among *M. tb* clinical isolates and appear to be transmitted at higher rates relative to mutations with high *in vitro* fitness costs (18, 23). Candidate loci involved in resistance and compensation for its fitness effects have been identified previously by screening for homoplastic variants (i.e. mutations that emerge more than once on the phylogeny) that are significantly associated with drug resistant phenotypes (24) and genes with higher *dN/dS* (ratio of non-synonymous versus synonymous mutations) in resistant compared to sensitive isolates (25). Application of these methods to whole genome sequence data from *M. tb* clinical isolates has recovered known drug resistance loci, as well as loci associated with cell surface lipids and biosynthesis, DNA replication, and metabolism.

The goal of the present study was to use patterns of genetic diversity at known drug resistance loci to identify the population genomic signatures of positive selection in natural populations of *M. tb.* Using whole genome sequence data from two populations for which phenotypic resistance data were available, we have identified several distinct signatures associated with these loci under selection. Based on these results, we propose methods of identifying loci under positive selection, including novel drug resistance loci, in clonal bacteria such as *M. tb.*

## Results

We inferred the phylogeny of 1161 *M. tb* isolates from Russia and South Africa (see Methods, Supplementary Table 1) using the approximate maximum likelihood method implemented in FastTree2 (Figure 1). The majority of the isolates belong to L2 (*n* = 667) and L4 (*n* = 481). *M. tb* nucleotide diversity was similar to previous estimates from a globally distributed sample (26). We identified lineage-specific patterns in overall diversity, with L4 having higher diversity than L2 (π_L2_: 3.6 × 10^−5^, π_L4_: 1.5 × 10^−4^). Previously published analyses of whole genome sequence data from L2 indicate that the majority of L2 isolates worldwide belong to a sub-lineage that has undergone relatively recent expansion (27, 28). In this sample from Russia and South Africa, the majority of L2 isolates belong to this sub-lineage, while the L4 isolates are associated with deeper branching sub-lineages. This likely contributes to the observed differences in diversity.

**Figure 1.**
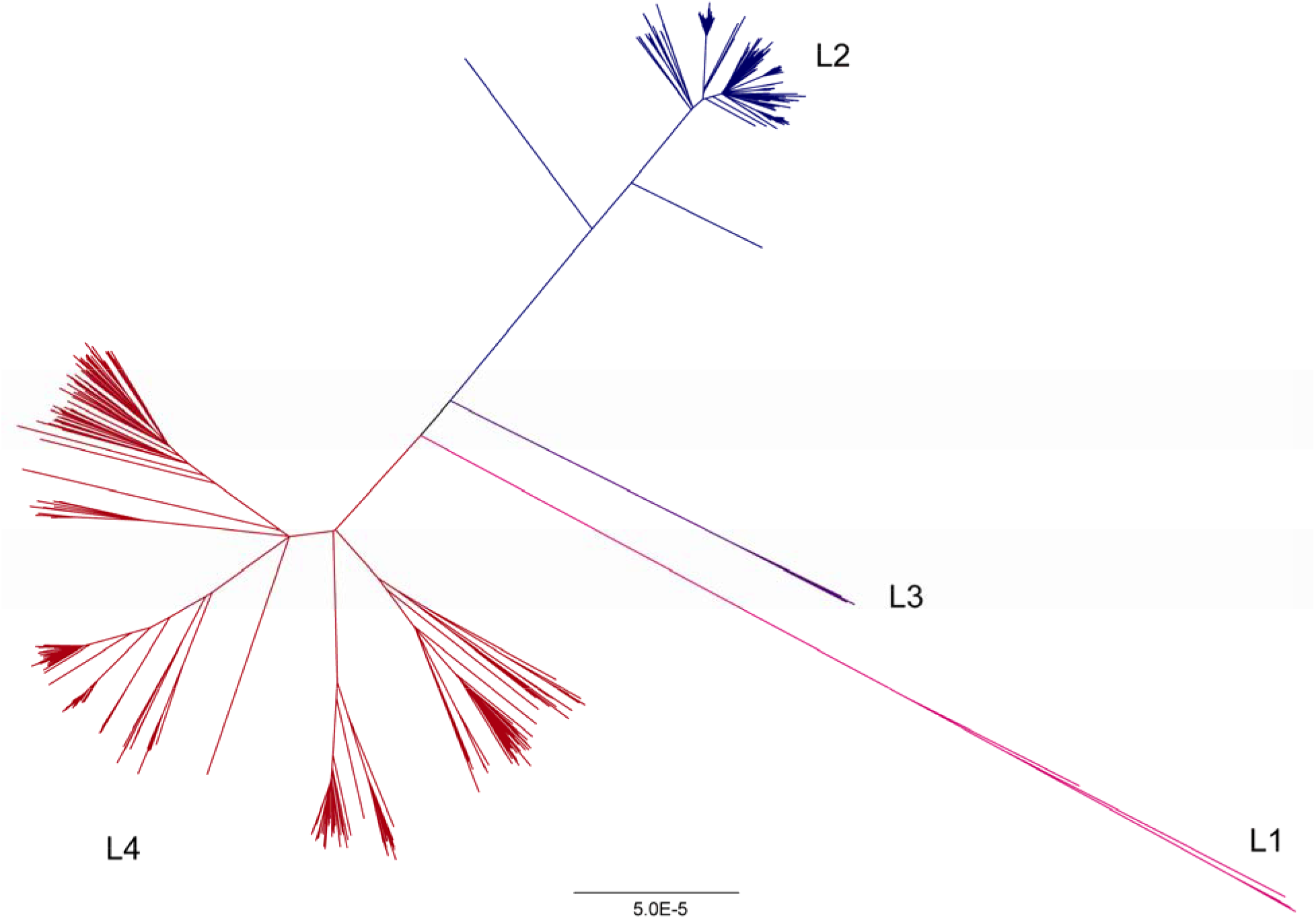
Phylogeny of *Mycobacterium tuberculosis* sample. The phylogeny was inferred using FastTree (60). Lineages are colored as follows: lineage 1 (L1) – pink, lineage 2 (L2) – blue, lineage 3 (L3) – purple, lineage 4 (L4) – red. Lineage 4 is associated with deeper branching sub-lineages in comparison with lineage 2.

Overall diversity of L2 was lower than L4 in our sample (Figure 2, *p* < 2.2 × 10^−16^). Seven hundred and sixty two of the isolates in our sample (66%) are resistant to one or more anti-tuberculosis drugs (Table 1).

**Figure 2.**
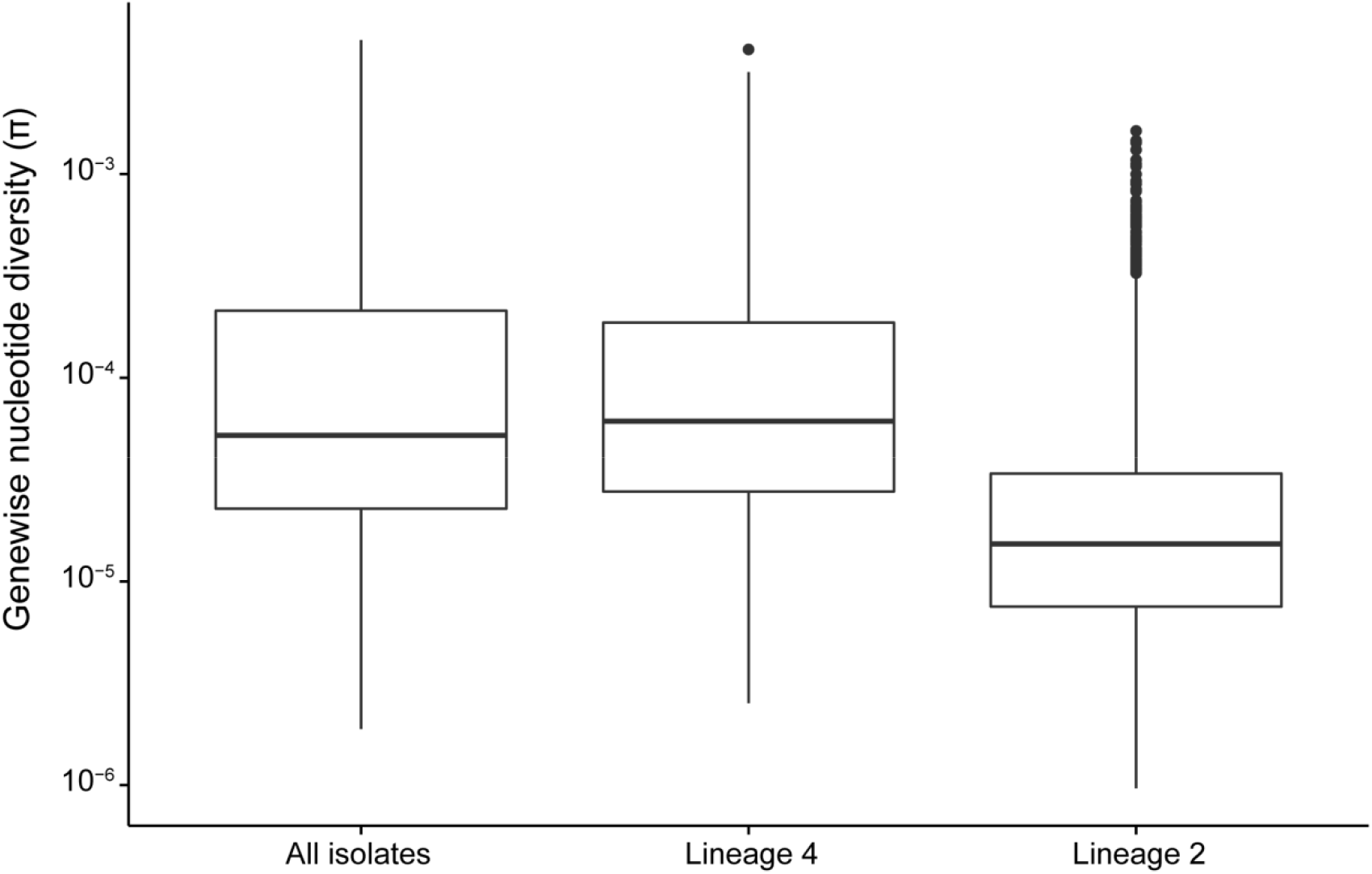
Distributions of gene-wise nucleotide diversity for all isolates, as well as lineages 4 and 2 considered separately. Repetitive regions of the alignment were masked. Sites were included in estimation of π if 95% of isolates in the alignment had a valid nucleotide at the position. We used Egglib to calculate statistics (64). Nucleotide diversity is lower in lineage 2 compared to lineage 4 (Welch Two Sample t-test, *p* < 2.2 × 10^−16^)

**Table 1.** Frequency of resistance in data set. AMI- amikacin, CAP- capreomycin, EMB- ethambutol, Et- ethionamide, INH- isoniazid, K- kanamycin, MOX- moxifloxacin, OFL- ofloxacin, PRO- protionamide, PZA- pyrazinamide, RIF- rifampin, STR- streptomycin.

| Drug | Resistant | Susceptible | Unknown |
| --- | --- | --- | --- |
| INH | 0.59 | 0.33 | 0.08 |
| STR | 0.53 | 0.39 | 0.07 |
| RIF | 0.50 | 0.43 | 0.07 |
| EMB | 0.30 | 0.59 | 0.11 |
| PZA | 0.21 | 0.67 | 0.12 |
| OFL | 0.16 | 0.39 | 0.45 |
| PRO | 0.15 | 0.22 | 0.62 |
| CAP | 0.10 | 0.41 | 0.49 |
| MOX | 0.09 | 0.41 | 0.49 |
| Et | 0.06 | 0.07 | 0.88 |
| AMI | 0.05 | 0.34 | 0.61 |
| K | 0.05 | 0.12 | 0.83 |

Drug resistant TB can be acquired as a result of *de novo* mutations within a patient or by infection with a resistant strain. When resistance is primarily mediated by *de novo* mutations, diversity should be similar in susceptible and resistant populations as resistance will arise on multiple genetic backgrounds. By contrast, if resistance develops primarily *via* transmission of resistant strains, the resistant sub-population should be less diverse than the susceptible sub-population. We compared the nucleotide diversity of susceptible and resistant sub-populations and found genome wide estimates of nucleotide diversity to be higher in isolates susceptible to a range of drugs for which phenotyping data were available (paired t-test, *p* = 0.029). In comparisons of gene-wise diversity in susceptible and resistant populations, we found that resistant isolates had a greater number of genes with no diversity, but levels of diversity within genes in which it was measurable were similar between resistant and susceptible populations (Figure 3).

**Figure 3.**
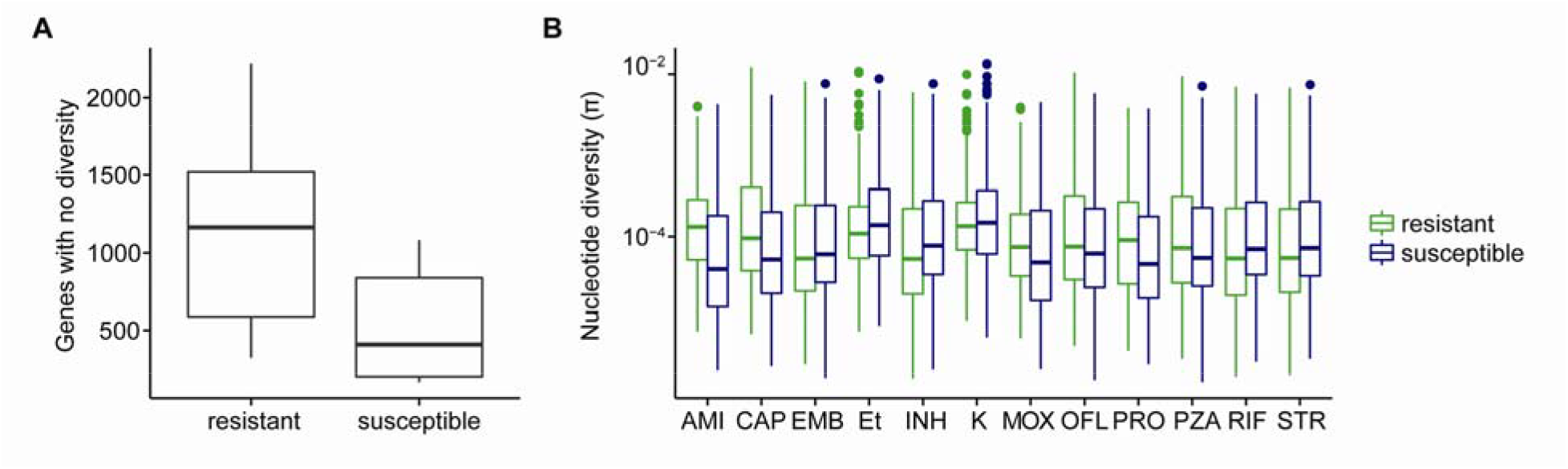
Diversity of resistant and susceptible isolates. A) Counts of genes with no nucleotide diversity in resistant and susceptible subpopulations. B) Genewise nucleotide diversity (excluding invariant genes) in susceptible and resistant isolates. Among genes in which it is measurable, nucleotide diversity is similar between resistant and susceptible isolates even when drug resistance associated genes and targets of independent mutation identified by Farhat et al. 2013 are removed (*p* = 0.13).

Of the 3,162 genes included in our analyses, 109 (3%) were invariant across all isolates in our sample. This is likely due to strong purifying selection on these genes. An additional 136 genes harbored variation in the drug susceptible populations but were invariant across all of the drug resistant populations. We did not observe the converse, i.e. genes that were invariant in susceptible isolates specifically, which supports the conclusion from genome wide diversity estimates that resistant isolates represent a subset of the diversity found in susceptible populations and suggests that there may be purifying selection that is specific to the setting of drug resistance. In order to evaluate whether the observed pattern was likely to arise by chance, we performed weighted random sampling of genes. The weighting was based on diversity in susceptible populations, assuming that genes with low diversity in susceptible populations are more likely to be invariant in resistant populations. After randomly sampling genes in each drug resistant population 1000 times, we found that no samples contained shared genes amongst all resistant populations (first and second line drugs). This suggests that specific genes tend to lose diversity in the setting of drug resistance, which could result from purifying selection specific to this setting. A potentially important caveat is that in our data set, drug resistant populations are not independent and the same isolates are often resistant to multiple drugs. Since resistance to second line drugs frequently arises on genetic backgrounds already resistant to one or more first line drugs, we repeated the sampling with only first line drugs and found that the maximum number of sampled genes shared across all populations was 2 (median 0). Overall, these results suggest that certain genes are more likely to lose diversity as drug resistance evolves, but we cannot completely rule out the possibility that the pattern arose as a result of overlapping membership in resistant populations.

We compared diversity of drug resistance associated genes (Table 2) with the rest of the genome using two measures of diversity: average pairwise differences (π) and number of segregating sites (θ). We found the resistance genes *gid, rpsL,* and *pncA* to be in the top 5^th^ percentile of gene-wise π and/or θ values. *rrs* and *ethA* are in the top 5^th^ percentile of θ, but not π. Surprisingly, despite being a target of multiple drug resistance mutations (Table 2), we did not identify extreme levels of diversity in *katG* (80^th^ and 82^nd^ percentile of π and θ, respectively).

**Table 2.**
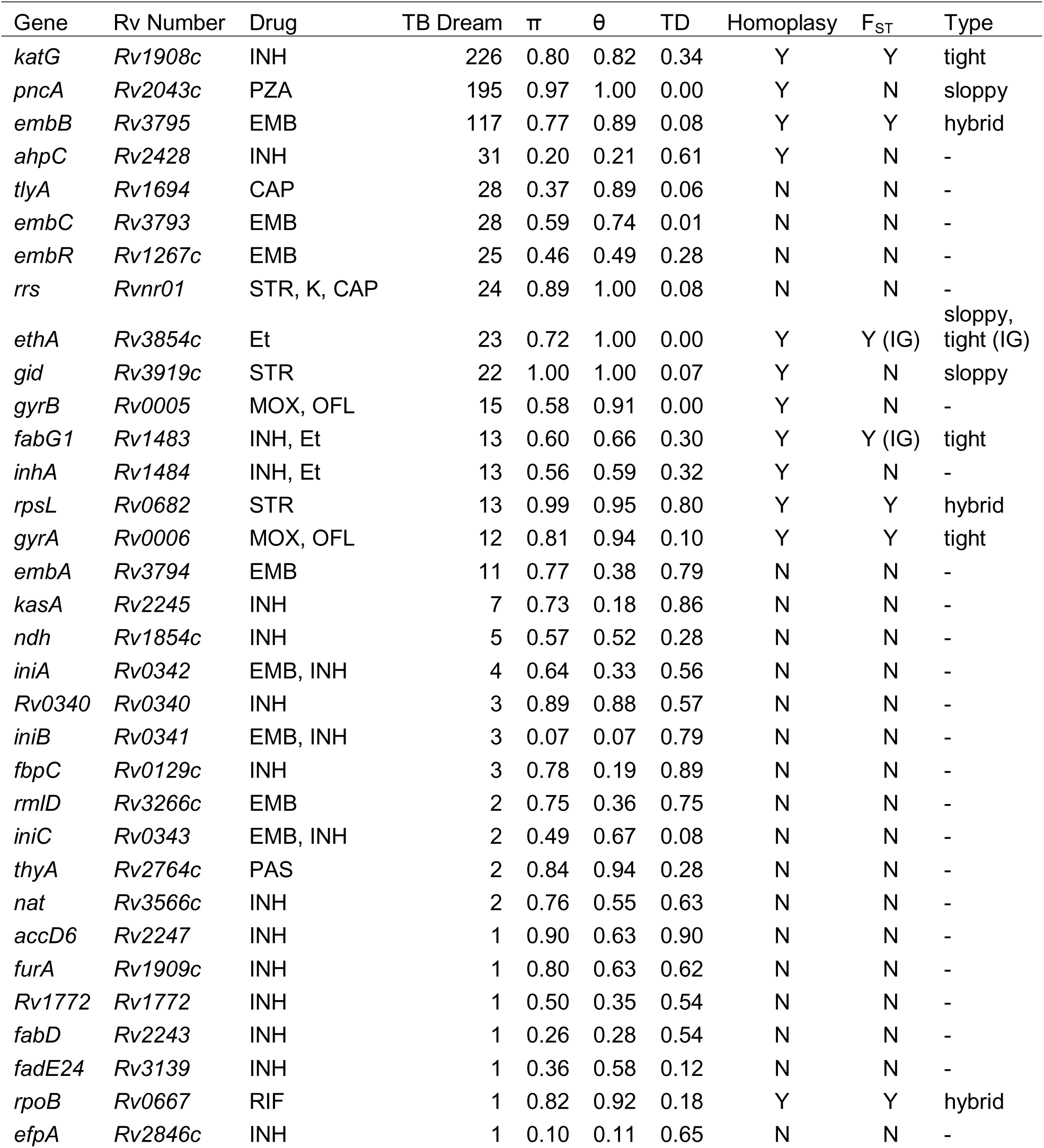
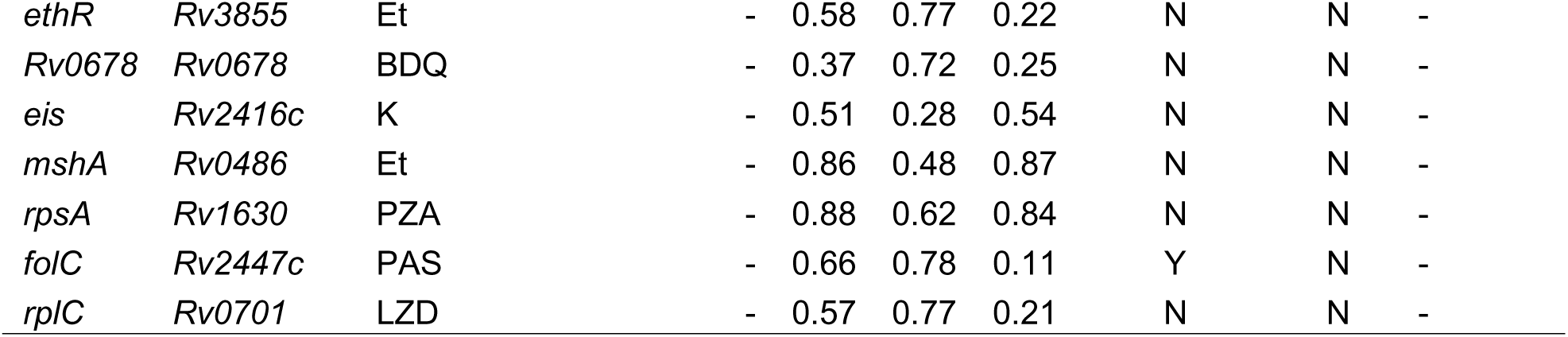
Signatures of selection in known drug resistance genes. The number of distinct entries in the TB Drug Resistance Mutation Database for each gene is reported in TB Dream column. π and θ are the percentiles for each diversity value, respectively. TD is the percentile of the residual after linear regression of Tajima’s D with gene length. Genes with homoplastic SNPs are indicated with ‘Y’ in the Homoplasy column. If a homoplastic SNP was also an F_ST_ outlier, it is indicated with a ‘Y’ in the F_ST_ column. Genes are classified as tight, sloppy, or hybrid targets of selection based on diversity, homoplasy, and F_ST_ results. (IG) indicates an intergenic SNP.

We also examined gene-wise diversity values within each lineage to look for lineage specific high diversity genes. In both L2 and L4, *gid, rpsL, pncA, ethA,* and *thyA* were in the top 5^th^ percentile of diversity (π and/or θ). In L2, *rpoB, embB, Rv1772,* and *folC* were additionally in the top 5^th^ percentile of gene-wise π and/or θ values. In L4, *Rv0340* was in the top 5^th^ percentile of gene-wise π and/or θ. While *rpoB* and *embB* were not in the top 5^th^ percentile of gene-wise θ in L4, they still had high diversity (91^st^ and 82^nd^ percentile, respectively). The lineage specific differences in diversity of *Rv1772, folC,* and *Rv0340* suggest that there are interactions between these loci and loci that differentiate L2 and L4.

We used gene-wise estimates of Tajima’s D to investigate gene specific skews in the site frequency spectrum that could result from selection, where negative values indicate an excess of rare variants and positive values indicate an excess of intermediate frequency variants. We previously identified a relationship between gene length and gene-wise estimates of Tajima’s D for *M. tb* (26), and this finding was corroborated here *(R^2^* = 0.3 after log_2_ transformation). In order to identify genes with extreme values of Tajima’s D - out of proportion with their length - we performed linear regression on log_2_ transformed gene lengths and Tajima’s D values and identified genes with the largest residuals (Figure 4). *pncA, ethA,* and *embC* all had Tajima’s D values lower than expected based on their length (5^th^ percentile of residual values). This indicates that these genes contain an excess of rare variants compared to other genes in the genome. Excess rare variants can result from a population expansion, a selective sweep, or purifying selection.

**Figure 4.**
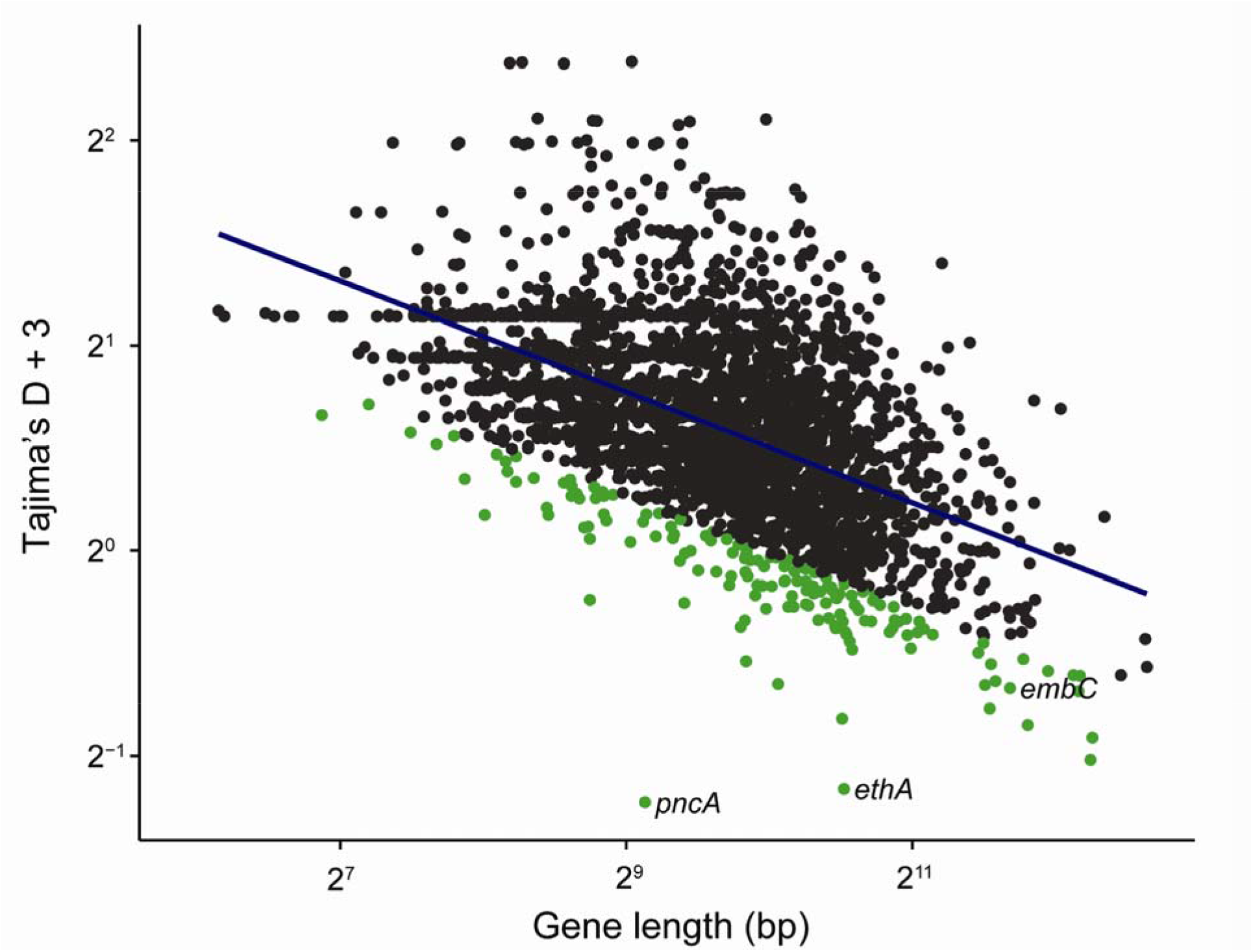
Gene-wise Tajima’s D and gene length. Repetitive regions of the alignment were masked. Gene lengths have been log transformed (base 2). We added a constant value (3) to all Tajima’s D values to make them positive and log transformed (base 2), as with the gene lengths. The linear regression line is plotted in blue. Genes with regression values in the lower 5% are highlighted in green. Drug resistance associated genes in this group are labelled. While negative Tajima’s D is normally associated with purifying selection or a recent selective sweep, we find that drug resistance genes with negative Tajima’s D also have high nucleotide diversity. We hypothesize that patterns of diversity at these genes have been affected by relaxation of purifying selection and positive selection in association with for drug resistance.

We calculated the ratio of π and θ of resistance associated genes in isolates susceptible and resistant to first line drugs and identified genes with markedly different diversities in resistant and susceptible sub-populations (Figure 5A). Among resistance genes in the top 5^th^ percentile of gene-wise π and θ overall, diversity of *pncA* and *ethA* is relatively high among resistant isolates, whereas diversity of *gid* is similar in resistant and susceptible populations. We also examined differences in this ratio between isolates in L2 and L4 (Figure 5B). *Rv1772* and *embR* were more diverse in resistant isolates in L2, and *kasA* and *tlyA* were more diverse in resistant isolates in L4.

**Figure 5.**
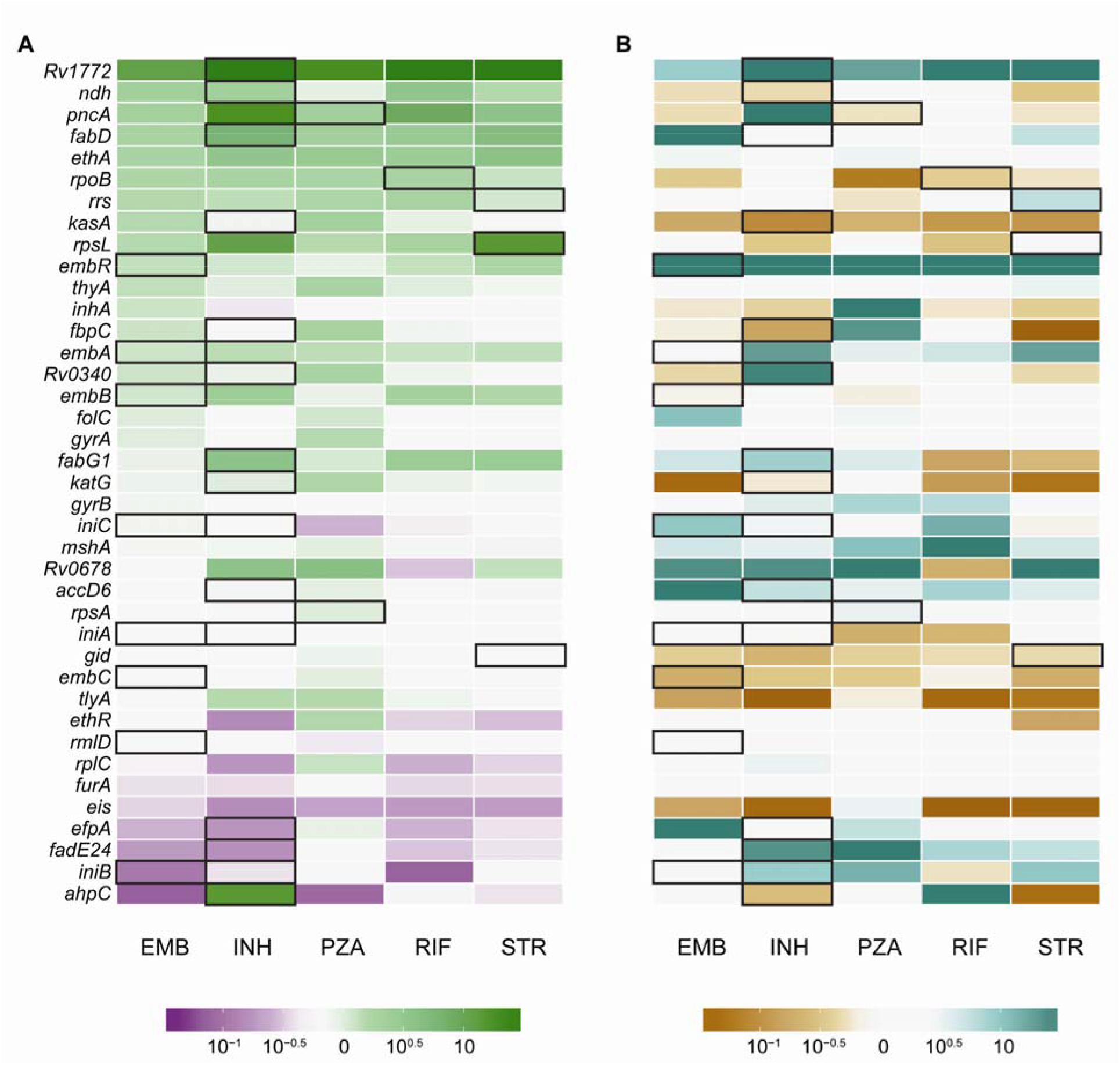
Ratios of nucleotide diversity in resistance associated genes. Genes with zero diversity were transformed to 1 × 10^−16^ before calculating ratios. Genes with ratios more extreme than 10^−1.5^ or 10^1.5^ are all filled with the deepest shade. Genes associated with resistance to each drug are outlined in black. A) Ratio of nucleotide diversity in resistant and susceptible isolates. Green genes are more diverse in resistant isolates, which could be due to diversifying selection and/or relaxation of purifying selection. Purple genes are more diverse in susceptible isolates, likely due to increased purifying selection. White genes have similar diversity in resistant and susceptible isolates. B) Comparison of ratios in lineage 2 and lineage 4. Teal genes are more diverse in lineage 2 resistant isolates, suggesting diversifying selection/relaxation of purifying selection specific to this lineage. Brown genes are more diverse in lineage 4 resistant isolates. White genes have similar diversity in lineages 2 and 4.

We used F_ST_ outlier analysis to identify single nucleotide polymorphisms (SNPs) and indels that exhibited extreme differences in frequency between susceptible and resistant populations. Our *a priori* expectation was that variants mediating resistance would be at markedly higher frequency in the drug resistant sub-population and that drug targets would be enriched among genes harboring variants with high F_ST_. After removing SNPs in regions corresponding to indels and variants at sites missing data for greater than 5% of isolates, the highest F_ST_ SNPs in comparisons of resistant and susceptible sub-populations to first line drugs are in *katG* (2155168, F_ST_ = 0.89, INH), *rpoB* (761155, F_ST_ = 0.72, RIF), and *rpsL* (781687, F_ST_ = 0.37, streptomycin (STR)). These SNPs were also F_ST_ outliers in the lineage specific analyses. We used a randomization procedure to assess the significance of observed F_ST_ values and found the maximum F_ST_ values after randomly assigning resistant and susceptible designations to be 0.023 for INH, 0.019 for RIF, and 0.018 for STR. In addition to SNPs within known drug resistance associated genes, we identified F_ST_ outliers in genes that may be novel targets for drug resistance (Table 3).

**Table 3.** Homoplastic F_ST_ outliers. Weir and Cockerham’s F_ST_ (wcFst) values in the top 1% of values genome wide are reported for each drug. For intergenic SNPs, the closest gene is listed. We identified mutations in genes previously associated with drug resistance (Known = Y) and novel putative resistance or compensatory mutations (Known = N).

| Location | Gene | Type | AMI | CAP | EMB | Et | INH | K | MOX | OFL | PRO | PZA | RIF | STR | Known | Lineage |
| --- | --- | --- | --- | --- | --- | --- | --- | --- | --- | --- | --- | --- | --- | --- | --- | --- |
| 1821 | *dnaN* | intergenic | - | - | - | 0.43 | - | 0.57 | - | 0.10 | - | - | - | - | N | all |
| 7570 | *gyrA* | missense | - | 0.33 | 0.11 | 0.46 | - | 0.66 | - | 0.29 | - | 0.18 | - | - | Y | all |
| 7572 | *gyrA* | missense | - | - | - | - | - | - | 0.06 | - | - | - | - | - | Y | all |
| 7581 | *gyrA* | missense | - | - | - | - | - | - | 0.07 | - | - | - | - | - | Y | all |
| 7582 | *gyrA* | missense | - | - | - | - | - | - | 0.35 | 0.22 | - | - | - | - | Y | all |
| 75233 | *icd2* | intergenic | - | - | - | - | - | - | 0.05 | - | - | - | - | - | N | all |
| 94388 | *hycQ* | synonymous | - | - | 0.07 | - | 0.12 | - | - | - | - | - | 0.12 | 0.13 | N | all |
| 230170 | *Rv0194* | missense | - | - | - | - | 0.12 | - | 0.05 | - | - | - | 0.12 | 0.13 | N | all |
| 332916 | *vapC25* | missense | - | - | 0.10 | - | - | - | - | 0.09 | - | 0.20 | - | - | N | all |
| 761155 | *rpoB* | missense | - | - | 0.31 | - | 0.58 | - | - | - | - | 0.10 | 0.72 | 0.41 | Y | all |
| 761161 | *rpoB* | missense | - | 0.33 | 0.09 | 0.51 | - | 0.71 | - | 0.13 | - | 0.16 | - | - | Y | all |
| 764817 | *rpoC* | missense | 0.19 | - | - | - | - | - | - | - | - | - | - | - | N | all |
| 781687 | *rpsL* | missense | - | - | 0.10 | - | 0.32 | - | - | - | - | 0.15 | - | 0.37 | Y | all |
| 922004 | *Rv0830* | missense | - | 0.30 | 0.12 | 0.43 | - | - | - | 0.10 | - | 0.21 | - | - | N | all |
| 1076880 | *Rv0965c* | synonymous | - | - | - | - | 0.12 | - | - | - | - | - | 0.12 | 0.13 | N | all |
| 1673425 | *fabG1* | intergenic | - | - | - | - | - | - | - | - | 0.11 | - | - | - | Y | all |
| 1673432 | *fabG1* | intergenic | - | - | - | 0.52 | - | 0.65 | - | - | - | - | - | - | Y | all |
| 1722228 | *pks5* | missense | - | - | 0.08 | - | 0.28 | - | - | 0.07 | - | 0.17 | - | 0.26 | N | all |
| 2122395 | *lldD2* | synonymous | - | - | - | - | - | - | 0.06 | - | - | - | - | - | N | all |
| 2155168 | *katG* | missense | - | - | 0.36 | - | 0.89 | - | - | 0.13 | - | 0.32 | 0.60 | 0.66 | Y | all |
| 2174216 | *Rv1922* | synonymous | - | - | - | - | - | - | - | - | - | 0.08 | - | - | N | all |
| 2207525 | *Rv1958c* | intergenic | - | - | - | - | - | - | - | - | - | 0.09 | - | - | N | all |
| 2422824 | *Rv2161c* | missense | - | 0.30 | - | 0.43 | - | 0.57 | - | 0.10 | - | - | - | - | N | all |
| 2660319 | *mbtF* | missense | - | - | 0.06 | - | - | - | - | - | - | - | - | - | N | all |
| 2715369 | *Rv3413c* | intergenic | 0.17 | - | 0.09 | - | 0.28 | - | - | - | - | - | 0.30 | 0.13 | N | all |
| 2866647 | *lppA* | synonymous | - | - | - | - | 0.12 | - | 0.07 | - | - | - | - | - | N | all |
| 2867298 | *lppB* | synonymous | - | - | - | - | 0.13 | - | - | - | - | - | - | - | N | all |
| 2867347 | *lppB* | synonymous | - | - | - | - | 0.13 | - | 0.06 | - | - | - | 0.12 | 0.14 | N | all |
| 2867756 | *lppB* | synonymous | - | - | - | - | 0.14 | - | - | - | - | - | - | - | N | all |
| 3500149 | *Rv3134c* | synonymous | - | - | - | - | - | - | - | - | - | - | 0.11 | - | N | all |
| 3550789 | *Rv3183* | synonymous | - | - | - | - | - | - | - | - | - | - | 0.12 | 0.13 | N | all |
| 3680932 | *lhr* | synonymous | - | - | - | - | 0.12 | - | - | - | - | - | 0.12 | 0.13 | N | all |
| 4001622 | *fadA6* | intergenic | - | - | - | - | - | - | - | - | - | - | 0.11 | - | N | all |
| 4247429 | *embB* | missense | - | - | 0.25 | 0.45 | 0.23 | - | 0.05 | 0.11 | - | 0.31 | 0.21 | 0.20 | Y | all |
| 4247574 | *embB* | synonymous | 0.19 | - | 0.07 | - | 0.27 | - | - | - | - | - | 0.30 | - | Y | all |
| 4327480 | *ethA* | intergenic | 0.20 | - | 0.07 | - | 0.27 | - | - | - | - | - | 0.30 | - | Y | all |
| 764948 | *rpoC* | missense | - | - | - | - | - | - | - | - | 0.06 | - | - | - | Y | L2 |
| 4248003 | *embB* | missense | - | 0.16 | - | - | - | - | - | - | - | - | - | - | Y | L2 |
| 698 | *dnaA* | missense | - | - | - | - | - | - | - | - | - | - | - | 0.10 | N | L4 |
| 60185 | *Rv0057* | missense | - | - | - | - | - | - | - | - | - | - | - | 0.06 | N | L4 |
| 761110 | *rpoB* | missense | 0.66 | - | - | - | - | - | - | - | - | - | - | - | Y | L4 |
| 764822 | *rpoC* | missense | - | - | - | - | - | - | - | - | - | - | - | 0.06 | Y | L4 |
| 781822 | *rpsL* | missense | - | - | - | - | 0.12 | - | - | - | - | - | 0.13 | 0.14 | Y | L4 |
| 2123145 | *lldD2* | missense | - | - | - | - | - | - | - | - | - | - | - | 0.06 | N | L4 |
| 2372550 | *dop* | missense | 0.64 | - | - | - | - | - | - | - | - | - | - | - | N | L4 |
| 2715344 | *Rv2413c* | intergenic | - | - | - | - | - | - | - | - | - | - | - | 0.06 | N | L4 |
| 2986827 | *Rv2670c* | missense | - | - | - | - | 0.16 | - | - | - | - | - | 0.17 | 0.15 | N | L4 |
| 4247431 | *embB* | missense | - | - | - | - | 0.11 | - | - | - | - | - | 0.11 | 0.07 | Y | L4 |
| 4248003 | *embB* | missense | - | - | - | - | - | - | - | - | - | - | - | 0.06 | Y | L4 |

Homoplastic SNPs – i.e. SNPs that evolve more than once on a phylogeny – are candidate loci under positive selection and have previously been used to identify resistance associated mutations in *M. tb* (24). Of the 235 genes with homoplastic SNPs that we identified in our sample, 13 are known to be associated with drug resistance (Figure 6), and resistance genes were significantly enriched among genes with homoplastic SNPs (Fisher’s Exact Test, *p* = 1.2 × 10^−4^). *pncA* had the largest number of homoplastic SNPs of any gene in the genome (n = 27 distinct SNPs that appear > 1 on the phylogeny). The SNPs identified in F_ST_ analysis were also identified as homoplastic (Figure 6). Our results suggest that complementary approaches based on homoplasy and F_ST_ outlier analysis can be used to identify SNPs associated with a trait of interest (in this case drug resistance). In addition to genic SNPs, we observed homoplastic SNPs that are also F_ST_ outliers in intergenic regions upstream of drug resistance associated genes (Table 3). These are candidate resistance and compensatory mutations with a regulatory mechanism of action.

**Figure 6.**
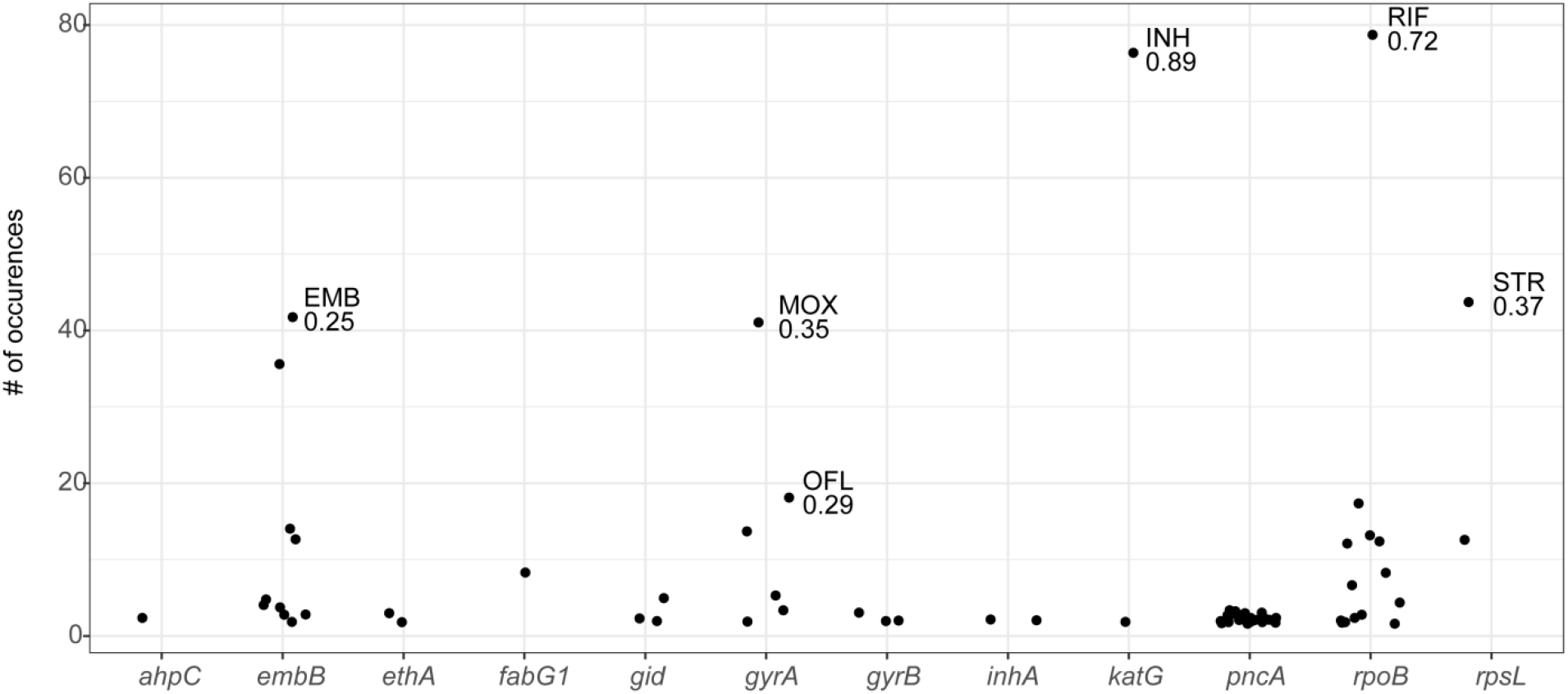
Homoplastic SNPs in drug resistance associated genes. SNPs with F_ST_ in the top 1% of genome-wide values are labeled with the population (associated drug resistance) and the F_ST_ value. *pncA* is remarkable for harboring diverse homoplastic mutations, each of which occurs relatively infrequently (“sloppy target”). *embB, gyrA, katG, rpoB* and *rpsL* harbor dominant mutations that occur frequently on the phylogeny and are strongly associated with resistant populations (“tight targets”).

In our analyses of indels, we controlled for the possibility that indels affecting the same gene may not be called in exactly the same position by considering indels within the same gene as identical. We identified four drug resistance associated genes with homoplastic indels: *gid, ethA, rpoB,* and *pncA.* F_ST_ values for the deletion in *gid* were in the top 5^th^ percentile for capreomycin (CAP), ethambutol (EMB), ethionamide (Et), kanamycin (K), ofloxacin (OFL), and pyrazinamide (PZA) resistant populations, but, interestingly, the deletion was not associated with STR resistance (F_ST_ = 0.04). Unlike homoplastic SNPs, homoplastic indels were not significantly enriched for drug resistance associated loci (*p* = 1).

We recovered 20 out of 40 known drug targets by identifying genes with extreme values of diversity, homoplastic SNPs, or SNPs that are F_ST_ outliers in comparisons of resistant and susceptible subpopulations. All genes with both extremely high diversity (top 5^th^ percentile) and homoplastic mutations were drug resistance associated (i.e. *gid, ethA, pncA,* and *rpsL).* We identified 67 genes with high diversity and Tajima’s D values more negative than expected based on gene length; only two of these were associated with drug resistance (i.e. *ethA* and *pncA).* Twenty out of 51 homoplastic SNPs that are also F_ST_ outliers fall within or upstream of known drug resistance associated genes. The remaining SNPs may be false positives or novel drug resistance mutations.

## Discussion

Highly virulent bacterial pathogens such as *M. tb, Yersinia pestis* (29), *Francisella tularensis* (30), and *Mycobacterium ulcerans* (31) appear to evolve clonally, i.e. with little to no evidence of lateral gene transfer. It is important to identify advantageous mutations in these and other organisms, as they are likely to be associated with phenotypes such as drug resistance, heightened transmissibility, or host adaptation. However, few methods are available for identifying loci under positive selection in the setting of clonal evolution. We adopted an empirical approach to this problem and used natural population data to characterize patterns of diversity at loci known to be under positive selection in *M. tb.*

In this analysis of clinical isolates from settings with endemic drug resistance, we found genome-wide diversity to be higher in susceptible *M. tb* sub-populations than in those resistant to first- and second- line drugs (with the exception of protionamide (PRO) and moxifloxacin (MOX) resistant populations). The observation of higher diversity in drug susceptible populations is consistent with a significant role for transmitted resistance in the propagation of drug resistant *M. tb.* A recent study of extensively drug resistant (XDR) *M. tb* infection in South Africa concluded that XDR cases result primarily from transmission of resistance, rather than *de novo* evolution of resistance mutations during infection (9). The primary studies for the sequence data analyzed here also identified clusters of drug resistant isolates (5, 6), suggesting that resistant isolates were being transmitted. Our results, along with these previously published observations, suggest that the fitness of drug resistant isolates can be high enough to allow them to circulate in endemic regions. As discussed below, the fitness effects of *M. tb* drug resistance mutations appear to vary substantially; the finding of transmitted resistance in this and other studies suggests that the fitness of isolates harboring low-cost mutations is comparable to that of susceptible *M. tb.* The populations in our study have a high burden of drug resistant TB, and the role of transmitted drug resistance may differ in other settings.

An alternative – but not mutually exclusive – explanation for the observation of higher diversity in susceptible populations is that drug resistant *M. tb* is under distinct evolutionary constraints that reduce average genome-wide levels of diversity. In support of this hypothesis, we identified a specific subset of genes that were invariant across drug resistant populations. Interestingly, while average diversity was lower for resistant sub-populations, the gene-wise diversity distributions had heavier tails, indicating there were more genes with extreme levels of diversity.

We found the genetic architecture of resistance to vary among targets, and resistance-associated genes tended to fall within categories that we term “sloppy”, “tight”, and “hybrid” targets of selection (the latter has a combination of tight and sloppy features and applies to *rpsL, embB,* and *rpoB).* “Sloppy” resistance genes are characterized by high levels of diversity. Genes associated with PZA, EMB, Et, and STR resistance (i.e. *pncA, gid, rpsL, rrs, ethA)* have high levels of diversity; some also had an excess of rare variants *(pncA, ethA, embC).* The finding that these genes accumulate multiple, individually rare mutations implies that there is a large target for resistance and/or compensatory mutations within the gene: that is, resistance can result from multiple different variants acting individually or in concert. In addition to its numerous rare mutations, *pncA* also contains the highest number of homoplastic SNPs (27 SNPs emerged more than once on the phylogeny) of any gene in the data set. Among the 62 non-synonymous *pncA* mutations in our dataset, 55 have been previously reported in association with drug resistance (TB Drug Resistance Mutation Database (32)). The newly described SNPs may mediate drug resistance or compensation for the fitness effects of other variants. Relaxed purifying selection is likely to play a role in concert with selection for diverse advantageous resistance mutations in the accumulation of diversity in *pncA* and other sloppy targets. The fact that numerous mutations are segregating in a natural population suggests that alterations to these genes are generally associated with negligible fitness costs. An *M. tb* strain harboring a deletion in *pncA* conferring resistance to PZA was estimated to be endemic in Quebec by 1800, long before the use of PZA for the treatment of TB (33–35). This supports the idea that purifying selection on *pncA* is relatively weak, which would contribute to its exceedingly high diversity and broaden the adaptive paths to resistance.

In contrast to *pncA, gid,* which is associated with low level STR resistance (36), does not appear to have the signatures of a “sloppy” target for resistance despite its high diversity. We identified just three homoplastic SNPs within *gid,* and previous studies have found that STR resistant isolates do not encode the same *gid* mutations (37). This could indicate that a multitude of mutations within *gid* confer resistance, but levels of diversity in the gene were similar in resistant and susceptible isolates. Previous studies of sequence polymorphism in *gid* have identified high diversity in this gene in both resistant and susceptible isolates (37–39): *gid* appears to be subject to relaxed purifying selection in the presence and absence of antibiotic pressure. Since *gid* mutations confer low level resistance, it’s also possible that mis-classification of resistance phenotypes contributed to the lack of differentiation we and others have observed between putatively STR resistant and susceptible sub-populations. In addition, mutations in *rpsL,* which cause high level resistance, could mask the contribution of *gid* to STR resistance.

We found some drug targets to be highly diverse in resistant sub-populations of either L2 or L4 (but not both), suggesting that resistance mutations in these genes interact with the genetic background; the fitness effects of mutations in these genes could, for example, vary on different genetic backgrounds. Lineage-specific F_ST_ outliers are another category of candidate locus with lineage dependent roles in drug resistance (Table 3). Epistatic interactions between drug resistance mutations and *M. tb* lineage have been reported previously: for example, specific mutations in the *inhA* promoter have been associated with the L1 and *M. africanum* genetic backgrounds (40, 41).

In contrast to “sloppy” targets, we discovered individual homoplastic SNPs associated with drug resistant sub-populations (i.e. with high F_ST_) representing “tight” targets of selection in genes conferring resistance to INH, RIF, and STR. Numerous resistance mutations have been described in *katG, rpoB, rpsL, embB,* and *gyrA,* but we find drug resistant sub-populations to be defined by a specific subset of mutations in these genes. This suggests that certain mutations are strongly favored relative to others conferring resistance to the same drugs when *M. tb* is in its natural environment. Antibiotic resistance can impose fitness costs on *M. tb* during *in vitro* growth, with the range of fitness costs varying among mutations, even within the same gene (18). Mutations can also have different fitness effects depending on the genetic background, but the most fit mutants were the same across *M. tb* lineages in a study of RIF resistance (18).

In our analyses, we found the dominant INH resistance mutation in *katG* to affect the serine at position 315. This change reduces affinity to INH but preserves catalase activity (42) and is associated with lower fitness costs than other *katG* mutants, both *in vitro* and in a mouse model (43, 44). This mutation was recently shown to precede mutations conferring resistance to other drugs during accumulation of resistance in evolution of multi-drug resistant *M. tb* (45). The dominant mutations we identified in *rpoB* (codon 450) and *rpsL* (codon 43) have also been found to have lower fitness costs *in vitro* compared to other mutations conferring resistance to RIF and STR in these genes (18, 44, 46). These results suggest that many of the findings regarding the relative fitness costs of *M. tb* resistance mutations *in vitro* and in animal models are relevant to the pathogen’s natural environment.

The fitness effects of mutations in *gyrA* (codon 94) and *embB* (codon 306) have not been measured; based on our homoplasy and F_ST_ results, we propose that they have lower fitness costs than other mutations in these genes and that they represent “tight” targets of selection. Mutations at *gyrA* codon 94 were previously found to be the most prevalent in a survey of *gyrA* and *gyrB* mutations in fluoroquinolone resistant *M. tb* clinical isolates (47). Interestingly, the mutation in *embB* codon 306 has been previously associated with acquisition of multiple resistances (48), and we find that this position is an F_ST_ outlier for all first line drugs in L4. This mutation is not an F_ST_ outlier in L2 (i.e top 5^th^ percentile), with percentiles for F_ST_ values ranging from 0.07–0.68 for first line drugs in this lineage. These observations suggest that the genetic background affects interactions among resistance mutations, and that *embB* 306 is important for acquisition of multidrug resistance in L4 but not L2.

We searched for indels with the signature of a “tight” target, i.e. homoplastic mutations segregating at markedly different frequencies in drug susceptible and resistant sub-populations. Unlike the pattern observed with SNPs, genes associated with drug resistance were not significantly enriched among those harboring homoplastic indels. We identified one homoplastic indel that was also an F_ST_ outlier - a deletion in *gid* that causes a frameshift. Patterns of variation in *gid* are complex and suggest a role for relaxation of purifying selection (i.e. in the accumulation of excess SNPs in both resistant and susceptible isolates) and perhaps a tight target associated with multi-resistance (i.e. this homoplastic/F_ST_ outlier deletion that was associated with resistance to CAP, EMB, Et, K, OFL, and PZA).

Our finding that, save for the frameshift mutation in *gid,* indels in resistance genes do not have the signature of “tight” targets suggests that they are generally associated with higher fitness costs than SNPs. Fifteen drug targets have been found in transposon mutagenesis experiments to be essential for *M. tb* growth *in vitro,* including *rpoB* and *rpsL;* deletions in these genes are likely to interrupt important functions (49). Deletions in non-essential genes could also have fitness costs. Deletions in *katG,* which is non-essential, can result in INH resistance but they are not observed as frequently in clinical isolates as the KatG S315 SNP, particularly among transmitted INH-resistant strains (23).

There are several limitations to our study. Resistance to multiple drugs was common in our sample, and in some cases it was difficult to identify patterns of diversity and population differentiation that were specific to individual drugs. Our results are also limited by the accuracy with which drug resistance phenotypes were determined and a lack of phenotypic data for some drugs (particularly second line drugs). Our sample was heavily skewed to lineages 2 and 4, and the results are not necessarily applicable to other *M. tb* lineages. Finally, the data analyzed here were generated with short sequencing read technologies, and we were thus limited to characterizing diversity in regions of the *M. tb* genome that can be resolved with these methods: regions that were masked from analysis (e.g. due to sequence repeats) may include unknown resistance targets. We also used an L4 genome (H37Rv) as a reference, and gene content specific to L2 may not have been identified.

We were not able to recover all drug resistance associated genes using the analyses performed here. This is likely a result of limited phenotypic data for some drugs and their associated targets (e.g. *thyA* and *folC,* which are associated with aminosalicylic acid (PAS) resistance). Our list of drug targets was dominated by genes associated with INH resistance, and signatures in genes that harbor rare resistance associated alleles may be subtle compared to the KatG S315 mutation found at high frequency in drug resistant populations.

We identified 31 SNPs that do not fall within the list of known drug resistance genes, which both emerged more than once on the phylogeny (homoplasies) and were segregating at markedly different frequencies in resistant and susceptible sub-populations (F_ST_ outliers). These SNPs may be novel resistance determinants; notably, all non-synonymous SNPs within this group are in genes linked with drug resistance in other studies (i.e. they are in genes encoding efflux pumps, genes differentially regulated in resistant isolates or in response to the presence of drug, potential drug targets, or genes in the same pathways as drug targets or resistance determinants) (50–54). In addition to a direct, previously unrecognized role in resistance, these SNPs could compensate for fitness costs of drug resistance. For example, we identified a homoplastic F_ST_ outlier in *rpoC,* and mutations in *rpoC* have been shown to compensate for RIF resistance in experimental evolution studies (22).

Intriguingly, we found lipid metabolism genes to be enriched in the list of genes harboring homoplastic SNPs (p = 0.013). We’ve previously shown that these genes have extreme values of diversity in a global sample of *M. tb* isolates and within individual hosts (26), suggesting that lipid metabolism genes may also be under positive selection in *M. tb* populations. The results presented here could be extended by phenotypic characterization of lipid profiles and identification of homoplastic variants that are at markedly different frequencies in isolates with distinct lipid profiles.

Here we have used drug resistance loci in *M. tb* to identify the signatures of positive selection in a clonal bacterium. We found these loci to be associated with distinct patterns of diversity that likely reflect differing genetic architectures underlying the traits under selection. The evolutionary path to resistance is broad for some drugs with “sloppy targets”, whereas for drugs with “tight targets” the means of acquiring resistance appear more limited. This is likely due to fitness effects of resistance mutations in *M. tb’s* natural environment, as numerous resistance mutations have been identified in tight target genes. We also found evidence suggesting that there are important interactions among loci during the evolution of resistance. Our results suggest that purifying selection on a subset of genes intensifies in the setting of resistance, which could reflect epistatic interactions and/or a response to the metabolic milieu imposed by antimycobacterial agents. The results presented here can be used to create more realistic models of resistance evolution in *M. tb* and to develop novel strategies of preventing or mitigating the acquisition of resistance. For example, the narrow path to resistance for drugs with tight targets reveals potentially exploitable vulnerabilities, as does the finding of interdependencies among specific loci and the genetic background in the evolution of resistance and multi-resistance. As new TB drugs become available for clinical use, the approach outlined here can be extended to investigate their architectures of resistance.

Efforts are underway to sequence and perform drug susceptibility testing on thousands of *M. tb* isolates with the goal of creating an exhaustive catalogue of drug resistance mutations and eventually using WGS to diagnose drug resistance in clinical settings (CRyPTIC project, <u>http://modmedmicro.nsms.ox.ac.uk/cryptic/</u>, last accessed: May 24, 2017). We found that loci under positive selection can be identified using relatively simple methods: “tight” targets are highly differentiated in their allele frequencies across phenotypic groups (i.e. F_ST_ outliers) and appear as homoplasies on the phylogeny; “sloppy” targets are characterized by high diversity and/or low Tajima’s D, as well as homoplasies. Extrapolating from patterns observed among known resistance variants, we have discovered new candidate regulatory and genic resistance variants. The methods used in this study are widely available and should scale to analysis of the large collections of genomic and phenotypic data that are currently being generated. This approach can be extended to identify novel resistance loci in bacteria for which drug susceptibility phenotypes are defined, as well as other positively selected loci in clonal bacterial populations.

## Methods

### Reference guided assembly

We downloaded sequencing read data from two large surveys of drug resistant *M. tb* in Russia (5) and South Africa (6). We used FastQC (55) and TrimGalore (56) for quality assessment and adaptor trimming of the reads. Trimmed reads were mapped to *M. tb* H37Rv (NC_000962.3) using BWA-MEM v 0.7.12 (57). We used Samtools v 1.2 (58) and Picard Tools (<u>https://broadinstitute.github.io/picard/</u>) for sorting, format conversion, and addition of read group information. Variants were identified using Pilon v 1.16 (59). A detailed description of the reference guided assembly pipeline is available at <u>https://github.com/pepperell-lab/RGAPepPipe</u>. We removed isolates with mean coverage less than 20X, isolates with percentage of the genome covered at 10X less than 90%, isolates where a majority of reads did not map to H37Rv, and isolates where greater than 10% of sites were unknown after mapping. The final data set contains 1161 *M. tb* isolates (Supplementary Table 1). The alignment was masked to remove repetitive regions including PE/PPE genes.

### Phylogenetic analysis

We estimated the approximately maximum likelihood phylogeny using the masked alignment from reference guided assembly with FastTree-2.1.9 (60). We compiled FastTree using the double precision option to accurately estimate branch lengths of closely related isolates. We used FigTree (http://tree.bio.ed.ac.uk/software/figtree/) for tree visualization.

### SNP annotation

A VCF of single nucleotide variants was created from the masked alignment using SNP-sites v 2.3.2 (61). SNPs were annotated using SnpEff v 4.1j (62) to identify synonymous, non-synonymous, and intergenic SNPs based on the annotation of *M. tb* H37Rv.

### Indel identification

Insertions and deletions were identified during variant calling with Pilon. We used Emu (63) to normalize indels across multiple isolates. We used a presence/absence matrix for the normalized indels for further analyses of indel diversity.

### Population genetics statistics

Whole genome and gene-wise diversity (π and θ) and neutrality (Tajima’s D) statistics were calculated using Egglib v 2.1.10 (64) for whole genome alignments and gene-wise alignments. Isolates were further divided by lineage and drug resistance phenotype. Sites with missing data due to indels or low quality base calls more than 5% of isolates in the alignment were not included in calculation of statistics. Values of Tajima’s D showed a correlation with gene length in our sample. To find genes with extreme values of Tajima’s D, we performed linear regression in R (65) on log transformed Tajima’s D values and gene length and identified genes with large residual values. To identify alleles with marked differences in frequency in resistant and susceptible isolates, Weir and Cockerham’s F_ST_ (66) was calculated using populations of resistant and susceptible isolates for each drug using vcflib v1.0.0-rc0-262-g50a3 (https://github.com/vcflib/vcflib). For non-biallelic SNPs, we calculated F_ST_ for the two most common variants.

### Homoplasy

We used TreeTime (67) to perform ancestral reconstruction and place SNPs and indels on the phylogeny. We identified homoplastic SNPs and indels as those arising multiple times on the phylogeny.

### Data availability

Unless otherwise noted, all data and scripts associated with this study are available at <u>https://github.com/pepperell-lab/mtbDrugResistance</u>.

## Acknowledgments

We thank members of the Pepperell Lab for their input on analyses and data visualization.

---

**Supplementary Table 1.** Accession numbers and lineage designation for sequence data passing quality control filters.

